# ITRAP2, a flexible and robust strategy to assign antigen recognition of T-cells in coupled single-cell TCR-pMHC assay

**DOI:** 10.64898/2025.12.19.695506

**Authors:** Grigorii Nos, Amalie Kai Bentzen, Marie Christine Viuff, Kristoffer Haurum Johansen, Jonas Birkelund Nilsson, Nikolaj Pagh Kristensen, Laura Stentoft Grand, Lasse Frank Voss, Mohammad Kadivar, Morten Nielsen, Sine Reker Hadrup

## Abstract

Determining T-cell specificity forms a crucial step toward understanding T-cell involvement in health and disease. Single-cell sequencing technologies allow for co-capture of TCR alpha and beta chains, and their antigen specificity can be determined through peptide-MHC (pMHC) multimer binding and capture of a co-attached barcode oligo. However, SC sequencing often includes a high level of dropouts and risk of cross-contamination. Similarly, barcoded pMHC readouts often suffer from significant background noise. These issues complicate the automatic assignment of pMHC recognition to TCR clonotypes. To overcome these challenges, we developed a method for data denoising - Improved T-cell Receptor Antigen Paring 2 (ITRAP2). This approach significantly reduces noise in single-cell pMHC readouts and allows for accurate identification of TCR specificity. ITRAP2 incorporates statistical tests and confidence metrics for each TCR-pMHC pairing, offering user flexibility in pairing rigor, and allows multiple pMHC assignments to the same T-cell clone in the event of cross-binding within the pMHC multimer library. We tested this method on an in-house generated dataset of 8141 single cells, screened for CD8⁺ T-cell binding using a panel of 100 different barcode-labelled pMHC multimers holding virus-derived peptides, and on a larger public dataset from 10x Genomics with 208,589 T-cells evaluated for recognition using a panel of 50 different pMHCs. In both datasets, ITRAP2 was able to recover TCR-pMHC hits that were missed either when investigating the raw data or analyzing the data using alternative tools. Importantly, we demonstrate that the size of the pMHC multimer library is crucial for accurate pMHC-TCR pairing and that a minimum of 25 pMHC multimer should be included to optimally determine background characteristic, and assigning true positive events.

## Introduction

CD8⁺ T-cells play a major role in the adaptive immune response by clearing cancerous and infected cells (Blum et al. 2013). Cells that display foreign peptides on their MHC-I molecules (pMHC) can be eliminated upon recognition by CD8⁺ T-cells (Gerlach et al. 2013; Farhood et al. 2019). CD8⁺ T-cells recognize the pMHC complex with their clonally derived T-cell receptor (TCR). Priming of CD8⁺ T-cells upon e.g. infection leads to the expansion of specific T-cell clones sharing the same TCR sequence, recognizing a peptide derived from the given infection or malignant tissue, and thereby enhancing the clearance of infected or cancerous cells presenting the given epitope (Gerlach et al. 2013).

Each TCR comprises an alpha (α) and a beta (β) chain, which together determine the T-cell’s specificity to a given pMHC (Davis and Bjorkman 1988). Since α and β chains are transcribed as separate mRNA molecules, pairing of these cannot be accomplished from bulk TCR sequencing, although attempts have been made to pair α-β chains based on their frequency. In contrast, single-cell TCR sequencing allows for direct pairing of α and β chains from individual T-cells, largely improving the utility of TCR data (Pai and Satpathy 2021; Maura et al. 2023). Therefore capture of TCRs from single-cell sequencing has emerged as the most efficient and least error-prone strategy to identify full-length TCR sequences (Han et al. 2014).

Understanding T-cell specificity has provided us with insights into T-cell biology and the role of T-cells in fighting infectious diseases and cancer. Recent technologies like barcoded pMHC multimer screening have generated large datasets dissecting CD8⁺ T-cell recognition to specific pMHC (Bentzen et al. 2016). However, bulk scale tools can only determine the presence of an antigen specific T-cell population, without revealing the number of pMHC-specific clones and their TCR sequence or T-cell state (Zhang et al. 2018).

Recent advancements in droplet-based single-cell capture technologies (Macosko et al. 2015; Klein et al. 2015) have enabled the coupling of multiple modalities at single-cell level. The field of T-cell biology has adapted parallel RNA-seq with TCR-seq, linking clonotypes with the T-cell phenotype. In the perfect scenario, a droplet captures a single cell into a gel-bead in emulsion (GEM), which labels each transcript with a GEM ID. Later, TCR sequences are amplified, with the capacity to trace α and β chain transcripts back to their cell of origin (Zemmour et al. 2018; Pai and Satpathy 2021).

Barcoded pMHC multimers have also been integrated into single-cell capture technologies, allowing pairing with other single-cell modalities, including TCR sequencing, providing an assay for pairing multiple TCR clonotypes with their pMHC recognition in parallel. However, achieving accurate pairing of single cell-derived TCR and pMHC recognition information has proven challenging due to the intrinsic noise from high dropout or multiplets rates and ambient contamination. Moreover, a substantial level of background noise among pMHC-derived barcodes is observed, likely caused by nonspecific cellular binding, making this data type difficult to handle (Povlsen et al. 2023; Zhang et al. 2021).

Large-scale TCR-pMHC dataset are highly warranted as input to build machine learning tools for pMHC assignment based on TCR sequence knowledge (Jensen and Nielsen 2024; Deleuran and Nielsen 2025; Croce et al. 2024), (Drost et al. 2025; Moris et al. 2021), and to understand similarity signatures observed across TCRs recognizing the same pMHC. It has also been demonstrated that a substantial fraction of pMHC-TCR pairs in the public domain are false, especially coming from single cell TCR-pMHC datasets (Messemaker et al. 2025). Such observations call for better and statistically robust strategies for TCR-pMHC assignment.

Several tools have been proposed to handle these challenges and “denoise” coupled single-cell TCR-seq with barcoded pMHC multimers data including ICON (Zhang et al. 2021) and ITRAP (hereafter referred to as ITRAP1) (Povlsen et al. 2023). However, while demonstrating promising performance, they both suffer from important shortcomings. For instance, ICON is dependent on internal negative controls used for signal normalization, and ITRAP1 is reliant on internally defined true positives used to define selection thresholds resulting for instance in an inability to identify cross-reactive TCRs.

To address these issues, we here present ITRAP2, a method that denoises single-cell pMHC readouts step-by-step by scaling each pMHC within the experiment and smoothing the pMHC values within each clone, using statistical tests for pMHC assignment, while maintaining user flexibility and clear graphical data representation. We applied this method to a newly generated single cell dataset with 8141 CD8⁺ T-cells screened for recognition towards 100 different pMHC multimers and a public dataset with 208,589 cells screened for recognition towards 50 different pMHC multimers (*Single Cell Immune Profiling Dataset by Cell Ranger v3*, n.d.-a). For both data sets, ITRAP2 was demonstrated to perform well, providing clear and easily interpretable outputs.

## Results

### Data description and noise sources identification

As an input to ITRAP2, we first generated an in-house dataset from a library of DNA-labelled pMHC multimers holding 100 different virus-derived peptides presented across 11 different HLA alleles. Epitopes were selected based on the prevalence of T-cell responses to common human viruses EBV-, CMV-, and influenza-derived epitopes, and 3 human self-peptides, established from previous work (Kristensen et al. 2024). This pMHC multimer library was used to screen for T-cell recognition in 11 donors, to ensure sufficient breadth of HLA alleles in the donor material. PBMCs were stained with DNA-labelled pMHC multimers with 10x 5’ v2-applicable single-stranded barcodes (see Methods) (Povlsen et al. 2023), TotalSeq-C hashtag antibodies, and fluorochrome-labelled lineage antibodies. Multimer-positive CD8⁺ T-cells were sorted and loaded for droplet-based single-cell capture using the 10x 5’ v2 platform. Consequently, we obtained both gene expression, scTCRseq and pMHC multimer binding information from each cell (Fig1A). The filtered dataset (for details see materials and methods) consisted of 8,141 single cells, of which 6,427 (79%) had at least one TCR β chain, and 4,487 (55%) had both productive chains available.

**Fig. 1:**
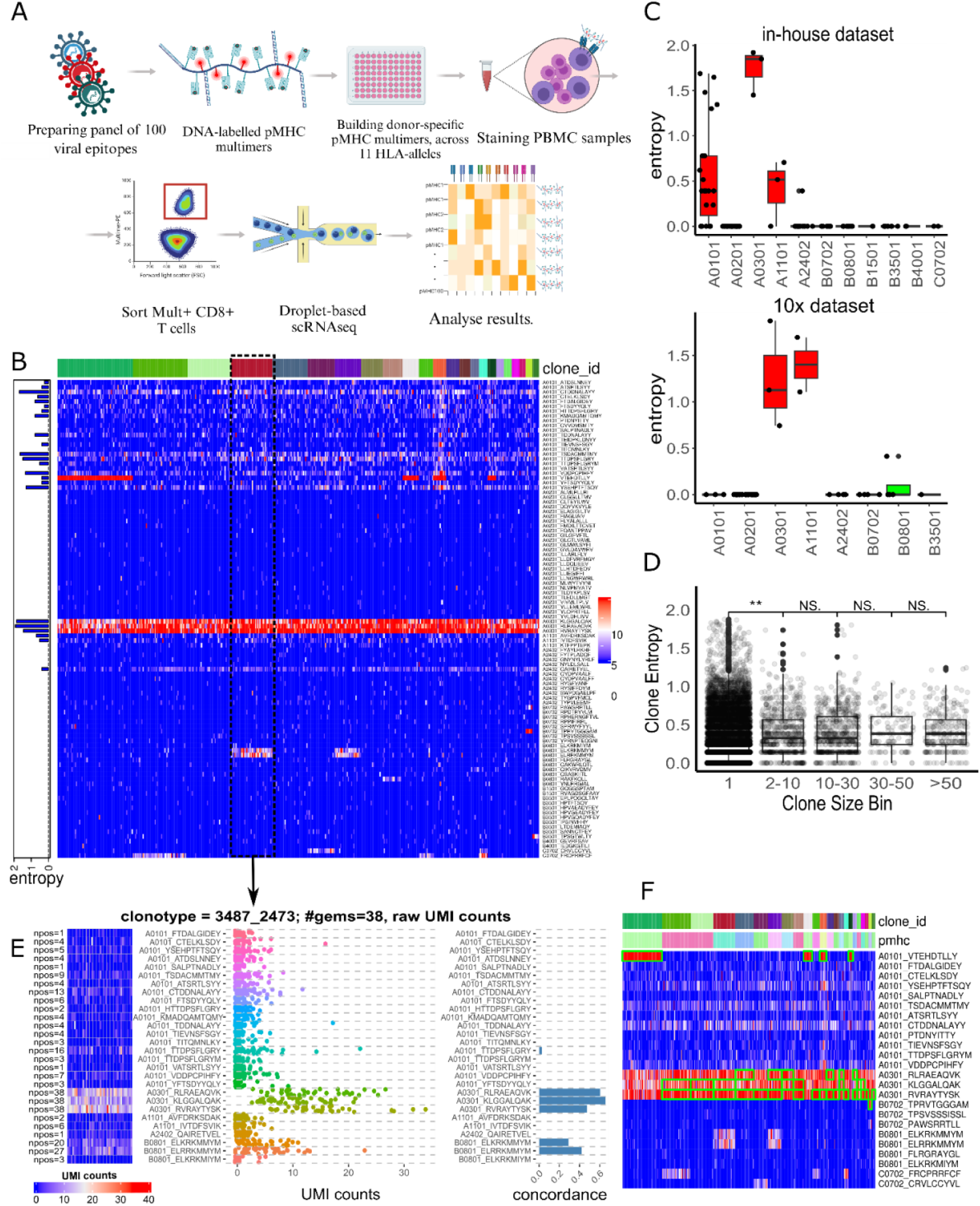
Experimental outline and initial look at raw pMHC UMI counts. **A)** An outline of the experimental process for generating single-cell TCR-pMHC coupled libraries. Created in BioRender. **B)** Heatmap of raw UMI counts for each pMHC barcode (rows) per GEM (columns) grouped by clonotypes. Noise level is measured in terms of entropy and displayed as a barplot on the left side of the heatmap for each pMHC. **C)** Boxplots presenting the entropy plotted for each pMHC, grouped per HLA allele for the in-house and 10x data sets. **D**) Boxplots with within-clone noise levels are measured by entropy, and grouped by clonotype size. Differences in clone size bins are estimated by Mann-Whitney test. **E)** The heatmap shows raw UMI counts of pMHC (rows) per GEM (columns) within a specific clonotype, with the number of gems with UMI > 0 marked on the left (npos). The middle dot plot shows UMI counts for each GEM of the given pMHC and the right barplot shows pMHC concordance, within this T-cell clone. **F)** Heatmap with UMI counts for selected informative pMHCs (rows) per GEM (columns) grouped by clonotypes. Green frames highlight pMHC specificities identified as significant using the outlier detection test within each clone, performed on raw UMI counts.

In addition to this in-house generated dataset, we analysed a public dataset generated by 10x genomics, consisting of 208,589 GEMs, of which 166,060 (80%) had at least one TCR β chain, and 145,213 (70%) had both productive chains available. This dataset was generated using pMHC multimers covering 44 common viral or tumour-associated T-cell epitopes and 6 negative control peptides across four donors. (*Single Cell Immune Profiling Dataset by Cell Ranger v3*, n.d.-b),

When examining the raw pmhc UMI counts (hereafter, “UMI counts” refer exclusively to pMHC barcode counts) (Fig1B, Sup Fig1A-D) grouped by TCR clonotypes, defined as the set of GEMs with identical α and β TCR sequences, we observed in both datasets substantial background noise, i.e. TCR clones with binding signals towards multiple divergent pMHCs, and specific pMHCs with binding signals dispersed across nearly every clonotype. To quantify the “noisiness” of individual pMHC, as a first measure, we used entropy; a metric from information theory that describes the level of uncertainty in the data. In short, for each clonotype, the entropy of a given pMHC is calculated using quantiles derived from the UMI counts of that pMHC across all GEMs. The overall pMHC entropy value is then obtained for each clone and then averaged across all clonotypes (see Materials and Methods). The underlying assumption is that less noisy pMHCs would yield UMI count values binned into a few quantiles within and across clonotypes, resulting in low entropy, whereas noisy pMHCs would have high quantile variation, leading to higher entropy values. We found that the entropy varied across HLAs, with HLA-A*03:01, HLA-A*01:01, and HLA-A*11:01 having the highest levels of entropy (median of 1.2 for HLA-A*03:01 and 1.4 for HLA-A*11:01), whereas pMHCs with no or limited noise had entropy values close to 0 (Fig1C). This variation in entropy driven by HLA was in part reproduced in the 10x dataset (Fig1C), indicating that certain HLAs are substantial sources of noise in both experiments (Fig1B,C). Additionally, we applied the same metric to quantify noise level, but per clone. We used the same pMHC-based quantile values, but with binned pseudobulk UMI count values within each clone, and summed over them to calculate entropy within a clone. Here, we expected noisy clones to have a broad range of quantiles present, whereas less noisy clones should mostly have the same low quantiles, with occasionally higher ones. From this analysis, we observed that clones of small size (i.e. formed by few or even single GEMs) generally had higher levels of noise than larger size clones (Fig1D). Since in most cases, the majority of T-cell clones in single-cell TCR datasets will be of small size, this underlines the additional challenge of assigning pMHC to TCR in clones of small size.

To quantify the consistency of the pMHC signal within a clone, we used a concordance metric (i.e. the fraction of GEMs in a clone that has a given pMHC as the highest UMI counts value among all pMHCs) as defined earlier (Povlsen et al. 2023). Although some clones showed a clear tendency toward a specific pMHC, the signal was often inconsistent, with several GEMs in a clone being both positive and negative outliers. Consequently, in raw UMI counts, concordance rarely reaches 100% (Fig1E) despite the assumption that GEMs within a given clone with identical TCR are expected to have a uniform signal towards the same pMHC(s). This variability suggests a substantial background noise or insufficient pMHC barcode capture and presents a challenge when attempting to assign pMHCs to a clone from the raw pMHC UMI counts. This can be illustrated when assigning the clonal pMHC specificities using methods for outlier detection based on, for instance, the raw UMI count values (in this case, the Rosner test (Rosner 1983), (see materials and methods) (Fig1F). In this figure, we first observe that the HLA-A*03:01 signal dominates every clone assignment, due to its intrinsic high UMI count values. We also visually observed multiple TCR-pMHC assignments with very inconsistent signals within clones, as well as some potentially false negative cases, where the outlier search missed visually clear specificities (Fig1F). Zooming in on an individual clonotype (Fig1E), despite having both a relatively high concordance towards HLA-B*08:01_ELRRKMMYM (42%) and HLA-B*08:01_ELKRKMMYM (25%) and visually convincing UMI counts patterns, this specificity was not assigned to the given clonotype due to the high background binding of HLA-A*03:01 multimers across all clonotypes (Fig1E). The issue is likely not unique to HLA-A*03:01, as we observed other HLA-A alleles having high entropy (Fig1C), so excluding all HLA-A*03:01 presented antigens would not be an optimal solution. These observations suggest that a procedure for signal normalization and denoising of the data is needed in order to make accurate TCR-pMHC assignments.

### Normalisation and Denoising

To effectively address the issue of background noise, we aimed to target the variations of the UMI count distributions both across the different pMHC and within a clone. First, to address difference across pMHCs, we implemented a modified version of mean (𝜇_𝑈𝑀𝐼_) centering and standard deviation (SD, 𝜎) scaling. This method iteratively removes UMI outliers for each pMHC before calculating the mean and standard deviation (for details refer to materials and methods), for z-score scaling (defined as 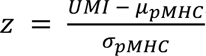). The outlier removal step was essential to avoid inflating the estimated mean and SD values. We showed the effect of this procedure, illustrating how the mean and in particular the SD values are highly inflated for some pMHC before applying the outlier removal (Suppl Fig2A). Similarly we illustrated how the outlier removal procedure supports the annotation of accurate TCR-pMHC assignments (Suppl Fig2B-C). Without outlier removal, a higher SD tends to lower the z-score, potentially compromising the detectability of positive signals, which are crucial for accurate TCR-pMHC assignment. The result of this is shown in Fig2A and B, where we observe how the modified scaling effectively normalizes UMI counts within each pMHC, removing the HLA-A*03:01 dominance while keeping potentially positive signals.

**Fig. 2:**
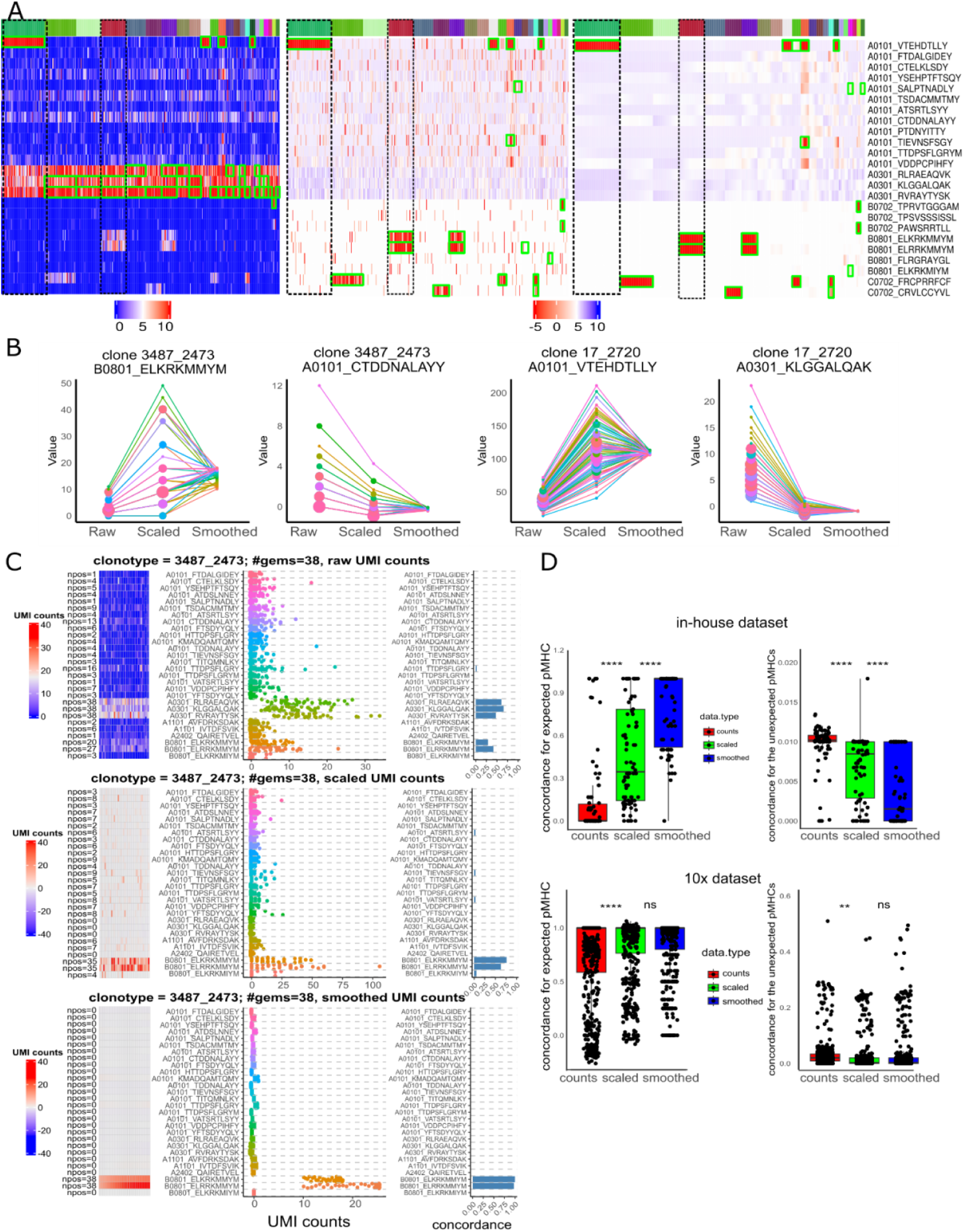
Denoising of pMHC UMI counts improves TCR-pMHC pairing. **A)** Heatmaps with raw (left), scaled (middle), and smoothed (right) UMI counts of pMHC (rows) per GEMs (columns) grouped by clones, TCR-pMHC pairings assigned in each step using Rosner outlier detection are shown as green frames. Black dotted box highlights clones 17_2720 (left) and 3487_2473 (centre) that are shown in later figures **B)** Dot plot showing changes of each GEMs UMI counts (y axis) within a clone from raw UMI counts, to scaled and LOESS smoothed states (x axis). **C)** Three plots showing the UMI counts distribution for clonotype 3487_2473 (same as shown in figure 1) for raw (top, identical to fig1E), scaled (middle) and smoothed UMI count values (bottom). **D)** Boxplots showing the pMHC concordance values (y axis) for each clone (each dot is a clone), based on raw, scaled and smoothed UMI count values, for assigned pMHC on the left and unassigned pMHC on the right. Mann-Whitney test was performed to compare concordance between data types.

Secondly, to address noise within a clonotype, we applied a LOESS (Locally Estimated Scatterplot Smoothing) regression to smooth the data and unify the signal (Cleveland and Devlin 1988). The smoothing works by fitting a localized regression model to subsets of the data, giving less weight to outliers, which effectively brings them closer to the major trend. In our approach, LOESS smoothing was useful for minimizing variability within each clonotype, ensuring that the pMHC counts reflect a consistent signal, as expected for T-cells with an identical TCR (Fig2A). We showed how smoothing effectively eliminates heavy outliers within the clones (Fig2B), making every GEM follow the same trend, thus, bringing low UMI values up within clones with a majority of high UMIs, and bringing unexpectedly high UMI counts down in clonotypes with a majority of low UMIs.

After the LOESS smoothing model is applied within each clone, we obtained a final “denoised” pMHC matrix. Next, to determine which pMHCs are most likely to bind to a given TCR, the denoised pMHC values for each GEM were aggregated into a pseudo-bulk for each clone. Subsequently, assigning a pMHC specificity from the panel became a task of positive outlier detection, i.e. identification the pMHC(s) that was significantly enriched from the total panel of pMHCs (the background distribution), and if no enrichment is observed, label clones as “non-assigned”. As a statistical outlier detection method the default choice was the Rosner test.

The Rosner test is a method for outlier detection in normally distributed data. We applied the method using the option to detect up to 10 outliers, since we expected a very limited number of different pMHC multimers to bind a given TCR clone. By applying the Rosner test, we could identify specific pMHC within clones if they were significantly enriched compared to the rest of the panel (Fig2A). This demonstrated an adequate assignment, with a consistent signal for each clone, matching the visually expected pMHCs paired with the given TCR (Fig2A,C).

Next, we observed that the denoised pMHC UMI counts, pseudobulked within a clonotype, followed an extreme distribution (Suppl. Fig3A). Therefore as an alternative, we could fit an extreme distribution for each clonotype and calculate the probability of highest denoised pMHC values to be outliers, given distribution parameters (see materials and methods).

To quantify the effect of each of the denoising steps, we calculated the pMHC concordance values within each clone for each step, using both the in-house and public 10x datasets (Fig2D). Here, we observed that each step significantly improves the consistency of the signal, since concordance for assigned pMHC per clone grows with each step, and decreases for unassigned pMHCs (Mann-Whitney test on in-house dataset - raw vs smoothed p=2.2e-16, 10x raw vs smoothed p = 2.2e-16). At the same time, we observed a decrease in average entropy for the noisiest HLA alleles (HLA-A*01:01, HLA-A*11:01, and HLA-A*03:01) in both datasets, when going from raw UMI counts to scaled and smoothed values (Suppl. Fig2B).

### Quality scoring for TCR-pMHC pairing

Even with the large effects of the denoising procedure described above, there were cases where noisy signal remained (Fig3A upper panel). Here, the assignments for donor #1 in the 10x dataset were shown after applying the scaling and smoothing denoising strategy, and within-clonotype outlier detection. This data set had three HLA-A*03:01 and two HLA-A*11:01 pMHC multimers that remained relatively noisy, and the outlier detection test ended up assigning specificity to these HLAs for almost half of the clones within this specific donor. This observation suggests that an additional step to reduce the noise within the pMHC data was needed.

**Fig. 3:**
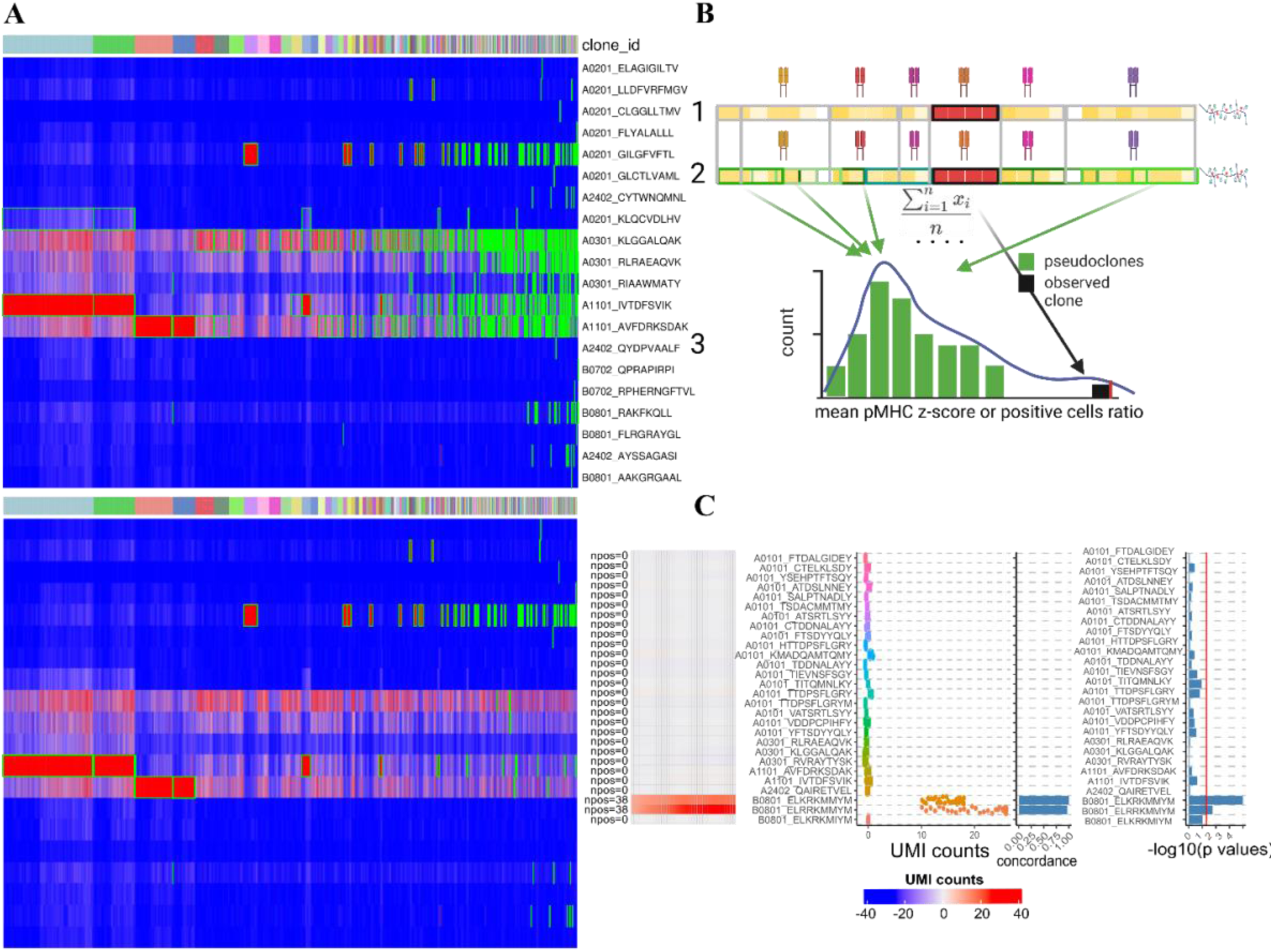
Applying the permutation test for the control of the noise within pMHC. **A)** Heatmap with UMI counts per pMHC (rows) per GEMs (columns) grouped by clonotypes for a set of selected specific clones from 10x dataset to showcase permutation test. Green frames depict assigned specificities without (top plot) and with (bottom) permutation test. **B)** Graphical representation of the permutation test. For each observed clone (black frame), pseudoclones are generated (different shades of green on a second row) of the same size, within the same pMHC space and the mean scaled UMI count value and a ratio of positive cells is calculated. The permutation test p-value is calculated from the proportion of pseudoclones that yield the same or greater metric than the observed clone. Created with Biorender. (credit: D.W.K., https://BioRender.com/z79j394). **C)** Heatmap of pMHC values from donor#1 in 10x dataset (identical to fig2C, lower plot), with the extra barplot on the right showing -10log p values of permutation test.

To address this, a permutation test for each pMHC-TCR pair was included to determine whether an observed clonotype exhibits higher and more consistent pMHC signal compared to simulated random clone. To make this test, we generated, for each assigned TCR-pMHC pair that passed the Rosner outlier detection test, N pseudoclones defined as a set of random GEMs, sampled outside the given clone and with random TCRs, but within the same pMHC space and with the same number of GEMs as the observed clonotype. For each pseudoclone, we then calculated two summary metrics; the mean scaled pMHC score and the ratio of positive GEMs (Fig3B, Suppl. Fig3D). Note, here we worked in scaled UMI space, and did not use smoothed pMHC UMI count values, since smoothing artificially elevates zero values and lowers positive outliers as well.

The null hypothesis assumed that pMHC signals within a clonotype are no stronger or more consistent than those obtained from random pseudoclones, whereas the alternative hypothesis stated that the observed clonotype exhibits higher and more stable pMHC UMI counts. Therefore the p-value was defined as the probability, under the null hypothesis, of generating a pseudoclone with a metric equal to or greater than that of the observed pMHC-TCR pair (Supp Fig. 3B). The two p-values derived from the metrics were then combined using Fisher’s method.

When applied as an additional filtering step, this permutation test effectively resolves the problematic over-assignments observed for HLA-A*03:01 and HLA-A*11:01, leaving only TCRs with extremely high and consistent signals for the HLA-A*11:01 pMHC (Fig3A lower panel). We also observed that the permutation test p-values appeared to be low where we expected, i.e. pMHCs with high concordance in a given clone, as we demonstrated in Figure 3C.

Applying this additional permutation test to the in-house and 10x data set, we showed that pMHCs with high entropy have higher median permutation test p-values, supporting that intrinsically noisier pMHCs on average produced lower-quality assignments (Suppl. Fig3C). Therefore, HLA-A*11:01, HLA-A*03:01 followed by HLA-A*01:01 all result in assignments with the highest p-values. At the same time, we can observe, across the HLAs with low entropy, that the specificity assignments coming from bigger clones are on average more robust, supported by the permutation test p-value, with the exeption of HLAs with high entropy, such as HLA-A*03:01 and HLA-A*11:01. (Suppl. Fig3E). Moreover the only big clones, that produced high p-value permutation tests, were the ones specific to HLA-A*03:01 and HLA-A*11:01 presented peptides.

Following this workflow, we additionally implement a measure of confidence for each pMHC-TCR pair. This metric takes into account the level of noise (pMHC entropy), size of the clone, pMHC concordance, and the delta gap value, or how distinct the outlier is in comparison with the background noise. To synthesize these parameters into a single metric, we developed a confidence score formula that users can adjust to balance sensitivity and precision trade-offs based on their specific requirements.

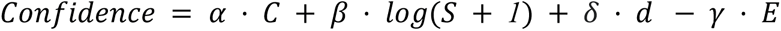

- 𝛼, 𝛽, 𝛾 — weights
- 𝐶 — clonotype concordance
- 𝑆 — clone size
- 𝐷 — difference between mean scaled pseudobulked pMHC UMI counts of specific mean pseudobulked background pMHC
- 𝐸 — pMHC entropy

The confidence score evaluates the reliability of the TCR-pMHC assignment within the dataset, and represents the effect size, while the permutation p-value assesses statistical significance against simulated controls and thus controls for the noise within a given pMHC. This distinction allows the user to potentially combine these metrics to tailor the TCR-pMHC pairing assignment to particular needs, driven by the biological context under examination.

### Validation of pMHC assignments using bulk T-cell recognition assessment

Due to the lack of TCR-pMHC-based ground truth data, we investigated to what degree the method could detect previously identified pMHC responses on a per-donor basis, in cases where we have an associated dataset with bulk pMHC T-cell screening results (Kristensen et al. 2024). Bulk pMHC screening can measure the presence of antigen-specific T-cells for a given pMHC within a sample (Bentzen et al. 2016). We correlated the observed bulk estimated number of pMHC-specific cells to the number of pMHC-specific GEMS identified by ITRAP2 (see Fig4A), and obtained a Pearson correlation coefficient of 0.72 (p=6.157e-10). Moreover, the few responses not captured in single-cell pMHC-TCR pairing, were all of low frequency in the bulk assay (below 1%) (Fig4B). This, together demonstrates strong correlation between the bulk and single-cell platform for identification of T cell recognition, with single-cell assay having slightly lower sensitivity.

**Fig. 4:**
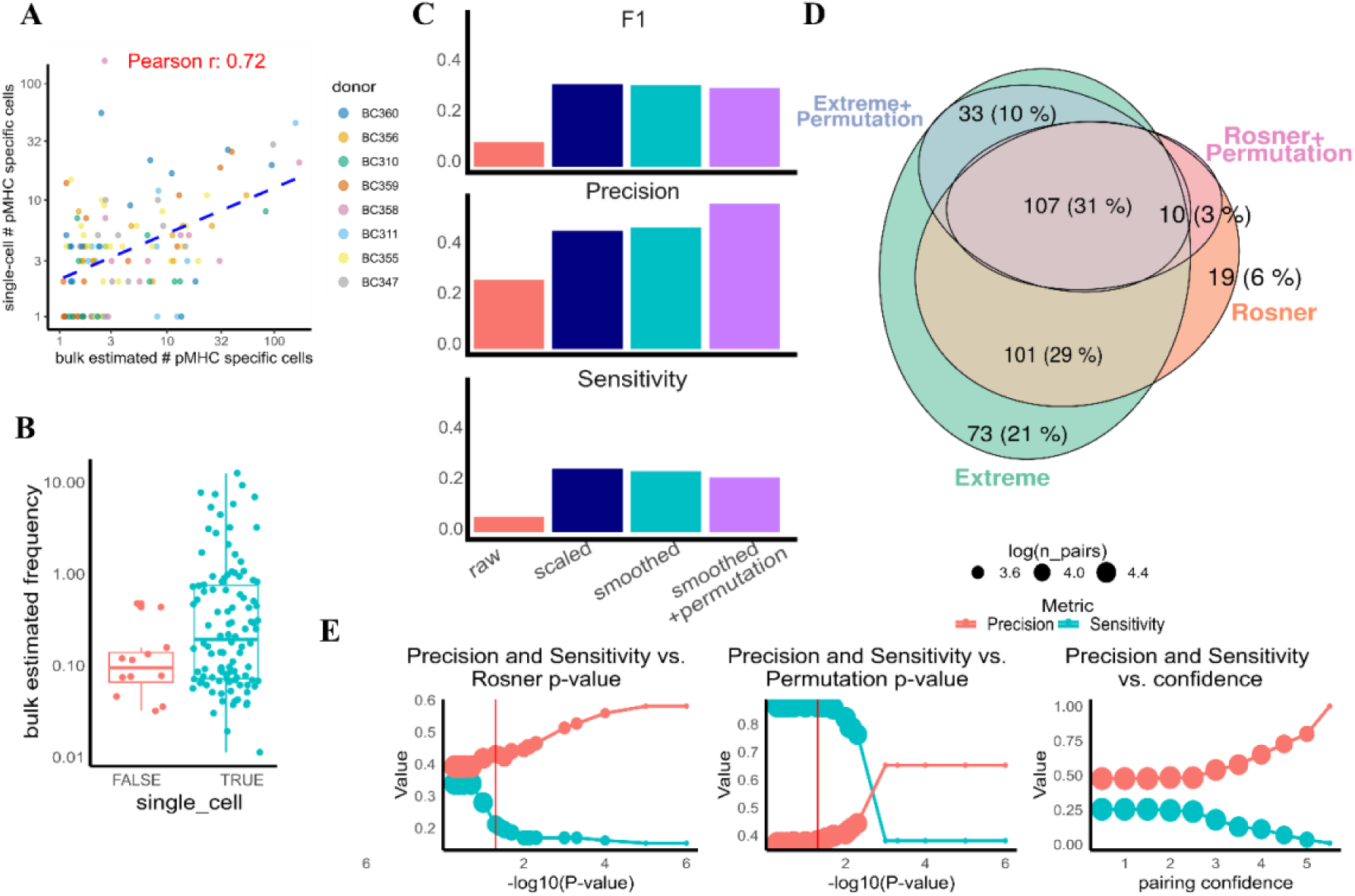
Quality control for TCR-pMHC assignments. **A)** Scatter plot with number of positive GEMs from single-cell data, across all clones on the y axis versus estimated number of cells for the T-cell population recognising the same pMHC, in the same donor, when assessed by bulk T-cell recognition analyses. **B)** Boxplot showing the estimated frequency of antigen-specific cells determined by the bulk screen, and split according to those identified in the single-cell data as well, and those not detected. Permutation test α of 0.05 is applied. **C)** Barplots with performance metrics changes per each ITRAP2 step (x axis), showing F1 scores (top), precision (middle) and sensitivity (bottom) **D)** Venn diagram, showing number of TCR-pMHC pairs identified with two different outlier detection methods, Extreme distribution and Rosner tests, with and without applying permutation test **E)** Scatter plots showing changes in precision and sensitivity with different p-value and confidence score cut-offs used.

We compared the pMHC specificities assigned in the in-house data using ITRAP2 with the antigen specificities previously determined in the same donor cohort using bulk pMHC screening (Kristensen et al. 2024). The performance was reported in terms of Precision, defined as a ratio of true positives to the overall number of assigned pMHC responses, and Sensitivity (or recall), defined as the ratio of true positives to a sum of true positives and false negatives (i.e. the total number of responses found in bulk screen), and F1-score as their harmonic mean. (Fig4C) From this analysis, we could observe that application of scaling, smoothing and permutation test all resulted in improved precision, hovewer permutation test resulted in minor loss of sensitivity.

Using the same metrics, we were also able to compare the performance of different within-clone TCR-pMHC assignment methods. A combination of Rosner and permutation tests or Extreme distribution with permutation test gave very similar performance (Fig4D), with high overlap in TCR-pMHC pairs’ outputs (78%, 107 out of 136 with permutation test).

We could further observe, as expected, that increasing the threshold for pairing TCR-pMHC based on permutation test p-value and confidence increases the precision of the pairing, though decreasing the sensitivity. (Fig4E).

### Optimal pMHC multimer panel size

The pMHC multimer assays are often custom-tailored, designed to address a certain research question. Technically, it is feasible to include as few as one pMHC in a given assay. However, since statistically determining the specificity of a clone essentially comes down to identifying outliers, we hypothesized that there is a minimal pMHC panel size necessary to enable accurate assignments.

To illustrate this, we iteratively, for each clone with at least one pMHC that passed the TCR-pMHC assignment (here based on the Rosner test), removed one pMHC from the library not identified as specific, and re-ran the ITRAP2 assignment pipeline on the reduced data set. In each iteration, we calculate the number of identified pMHC hits and record the number of incorrect assignments (assignments that differ from the assignments obtained using the original complete data set). In both datasets, inaccurate assignments started to accumulate when fewer than 20-30 pMHCs remained in the panel, (Fig5A). Similarly, we observed a gradual loss in the total number of assigned TCR-pMHC pairs using simulated panels with fewer pMHC (Fig5B). At the same time, the Rosner test’s p-value within the same TCR-pMHC pair on average got higher with a smaller panel size (Fig5C). In conclusion, these results suggested that having a panel size of at least 25 pMHCs is needed to ensure reliable TCR-pMHC pairing, because the approach starts to accumulate false-negative assignments around a panel of this size.

**Fig. 5:**
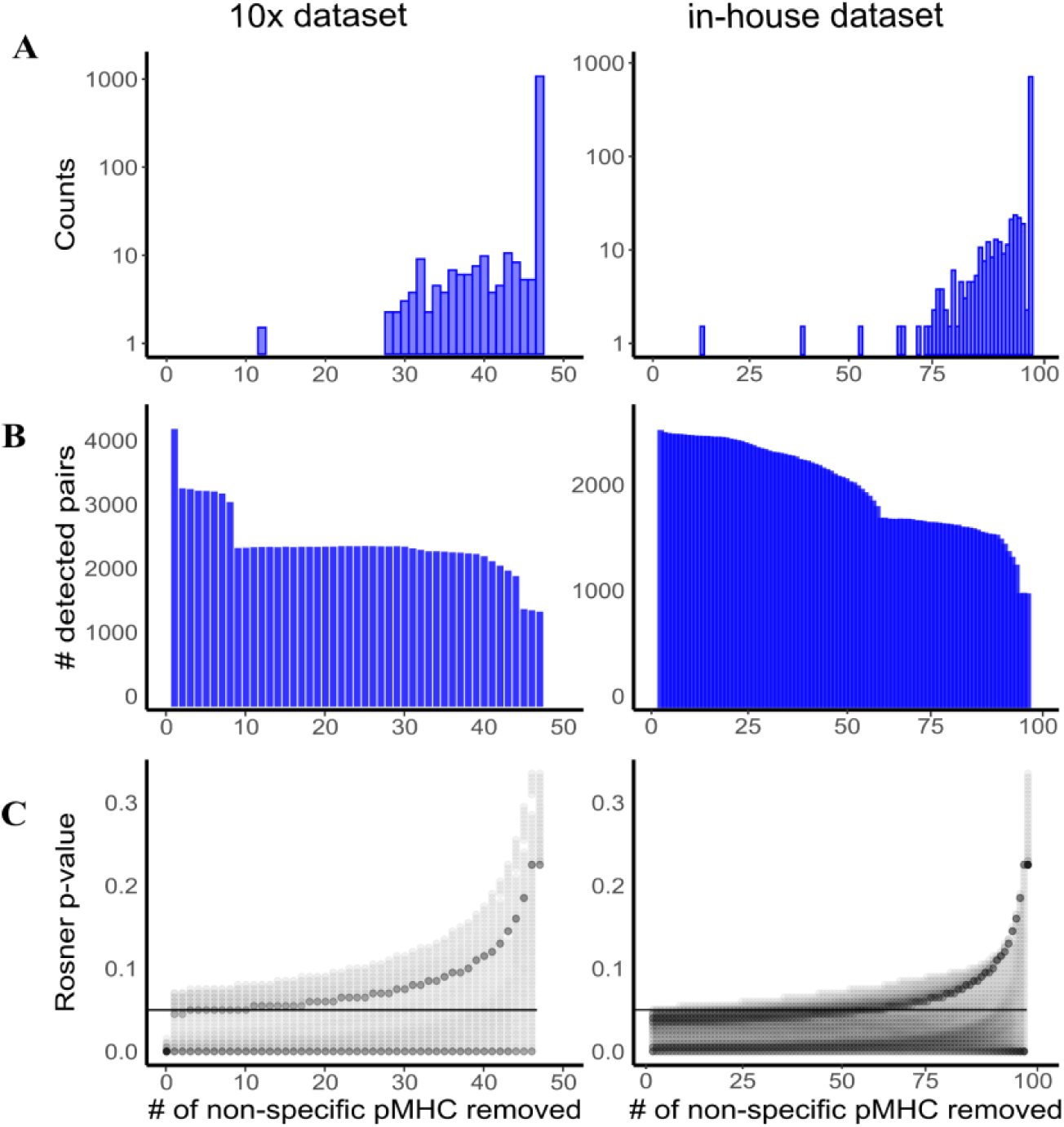
Optimal pMHC panel size. **A)** Barplots showing the number of wrong assignments (y axis) per the remaining number of pMHC in the panel (x-axis) if 1 random non-assigned pMHC is removed, until the test fail to perform. **B)** Barplots showing the number detected pMHC pairs (y axis) per the remaining number of pMHC in the panel (x-axis) if 1 random non-assigned pMHC is removed, until the test fail to perform. **C)** Dotplot showing Rosner test p values (y axis) per the remaining number of pMHC in the panel (x-asix) if random non-assigned pMHC is removed, until the test fail to perform. The darker color depicts the density of the dots in the coordinate

### Comparison with previous methods

As a final investigation, we compared TCR-pMHC assignments in the 10x dataset using the three methods, ICON, ITRAP1, and ITRAP2. Here, we could observe that the methods exhibit a rather low overlap, and especially for noisy pMHCs, the overlap is very small and even zero (Suppl. Fig4A). In contrast, for some well-known and low entropy pMHCs a high level of agreement between the methods was observed. At the same time filtering clones that are at least of the size of 3 GEMs also significantly increased the overlap (Suppl. Fig4B,5B). We observed a trend that relative overlap between TCR-pMHC assignments correlated negatively with pMHC noisiness, defined by entropy. (Suppl. Fig4D, E, 5B), which further suggested low reliability of high entropy pMHCs.

In Suppl. Fig4C, we show that ITRAP1 and ICON excessively assign specificities coming from high entropy HLA-A*03:01 and HLA-A*11:01 multimers. The assignments based on the ITRAP2 approach are visually more consistent with the pattern of high and stable UMI values within a clone (Suppl. Fig3A, Suppl. Fig4C).

We next investigated the distribution of pMHC UMI counts for clonotypes, where these three methods disagree. First, for each donor in the 10x data set, we took the all the clones that were assigned to a specific pMHC by at least one of the methods and plotted heatmaps with highlighted assignments by the three methods (Suppl. Fig4C). These visualisations illustrate that ITRAP2 is more consistent in TCR-pMHC assignments related to pMHC UMI signal within the clone, than ICON and ITRAP1.

To further compare ITRAP2 to ITRAP1, we ran ITRAP1 on our in-house dataset to compare performance. Due to the absence of a scaling procedure in the ITRAP1 method, it ends up assigning almost every single clonotype to HLA-A*03:01 presented peptides given their inflated and noisy UMI counts. If we, as a means to resolve this, removed HLA-A*03:01, peptides from the analysis, the second most noisy HLA, HLA-A*01:01 took over. This emphasised the importance of UMI counts for standardisation before applying specificity assignment to TCRs and showed the advantage of ITRAP2 over ITRAP1.

To further compare the performance of the different methods, we investigated the inter-pMHC versus intra-pMHC TCR similarity as proposed earlier (Povlsen et al. 2023). Reasoning that TCRs recognizing the same pMHCs (intra-specificity) should exhibit higher sequence similarity compared to TCRs recognizing other pMHCs (inter-specificity). Suppl. Fig5A showed the result of this evaluation in terms of the difference between the intra and inter-similarities for the subset of 25 pMHCs annotated with two or more TCRs for all three methods. As expected, the delta similarity metric is positive for most peptides for all three methods (16 of 25 for ITRAP1 and ICON and 18 of 25 for ITRAP2). The average delta value for the most represented 25 peptides, used on 10x dataset, is slightly higher for ITRAP2 compared to ITRAP1 and ICON. This result suggests that ITRAP2 performs at least on par with ICON and ITRAP1 using TCR similarity as the evaluation parameter. For the majority of the peptides we identified more than one specific TCR, keeping at least the same delta-similarity score. There are few peptides with lower number of detected TCRs, such as IVTDFSVIK, AVFDRKSDAK, GILGFVFTL, RAKFKQLL, but ITRAP2 outperformed ITRAP1 and ICON in terms of delta similarity metric, for these single peptides (Suppl. Fig5A).

We also examined three well-known epitopes, GILGFVFTL, RAKFKQLL, and ELAGIGILTV, where we expected a large number of T-cell clones. We visualized every clone that ICON, ITRAP1, and ITRAP2 assigned to at least one pMHC and plotted each clone set on separate heatmaps to investigate the TCR-pMHC assignments made by each method (Suppl. Fig5C). Here, we observed high UMI counts in almost every ITRAP2 assignment. In contrast, some low UMI counts in ICON and ITRAP1-assigned clonotypes is observed, again potentially indicating that these tools result in false positive assignments.

All together, we believe these results support that ITRAP2 is outperforming both ITRAP1 and ICON, providing a revised, user flexible and statistically driven strategy for accurate assignment of TCR to pMHC specificity in single-cell data.

### ITRAP2 usage

Based on the results described above, we implement the rescaling, smoothing, and outlier detection methods in an R package called ITRAP2. Below, we describe in detail the functionality of this tool.

The input to the method is a general 10x cellranger output, with count matrix and per-GEM (filtered_feature_barcode_matrix folder) annotations of TCRs, as CellRanger VDJ output (filtered_contig_annotation.csv) (Fig6A). Optionally, TCR output from CellRanger multi can further be processed with downstream VDJ tool of user’s choice, such as scRepertoir (Yang et al. 2025), Immunarch (Popov et al. 2025), Dandellion (Borcherding et al. 2020) or Changeo-o (Gupta et al. 2015). Similar datasets can be generated through other platforms, e.g. BD Rhapsody, and is expected to perform similarly, but has not been tested. If gene expression information is available, we recommend applying regular gene expression-based filters in your toolkit of choice, for removal of doublets (both based on hash-taging and GEX information) and dead/dying cells, as we expect bad quality GEMs to produce extra noise. We recommend using the Seurat toolkit (Hao et al. 2024) to save your pMHC matrix in an extra assay of your Seurat object and add clonotype annotation to the meta data. Next, we apply outlier-less scaling (function ScaleDataNoOutliers), smoothing (function smooth_pmhc), and TCR-pMHC pairing with the assign_pmhc functions. At this stage permutation test and confidence scoreare also calculated for each TCR-pMHC pair, that passed Rosner test. (Fig6B). In Fig6C we show the expected changes of the pMHC matrix with the abovementioned steps, the user will be able to track them, highlighting clonotypes with pMHC specificity. The visualisation of this clonotype-sorted pMHC UMI counts matrix is available in our package with a function called pmhc_heatmap().

**Fig. 6:**
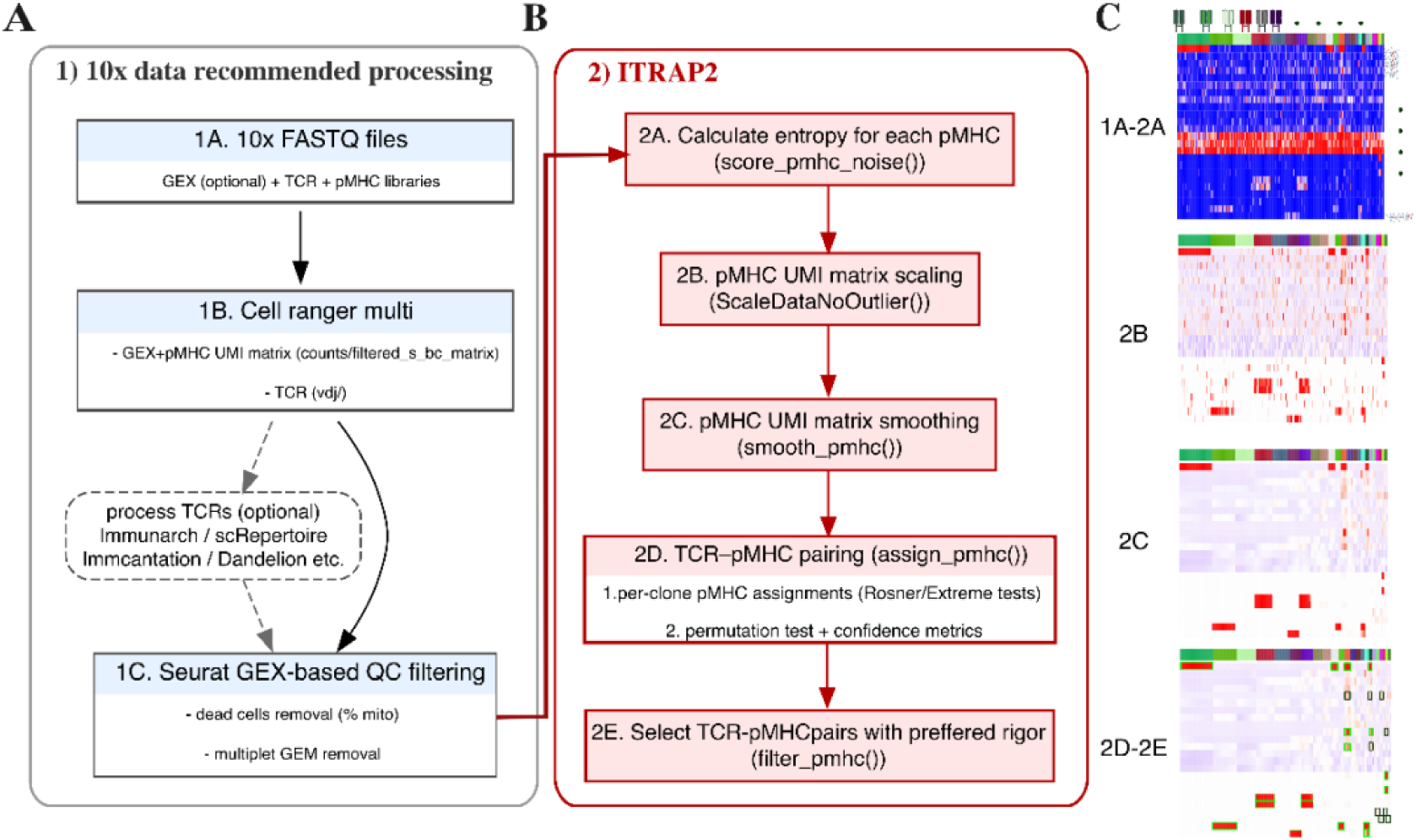
ITRAP2 usage. **A)** Flowchart representation for a recommended data processing prior to running ITRAP2. **B)** Flowchart representation of major steps in ITRAP2 **C)** Heatmap with GEMs, grouped as clonotypes (columns) and pMHCs (rows) colored by UMU counts in each step in ITRAP2. Steps from 1A-2A generate raw pMHC UMI count matrix, in step 2B scaled matrix is generated, in 2C per-clone smoothed and in step 2D we can assign pMHCs to TCRs (green and black frames) and afterwards permutation test can filter out lower confidence assignments (black frames)

As demonstrated in Fig1D, smaller clonotypes showed less clear signals, making pMHC-TCR pairing particularly challenging. Smoothing techniques are less effective for these smaller clones (and it is not feasible at all for single GEM clones) which limits the denoising options. However, we still provided users with the option to assign TCR-pMHC pairs for single-GEM clones using the same combination of the Rosner or Extreme distribution tests and the across-pMHC permutation test. We still observed cases of one GEM clones having a strong signal in some pMHCs with a low p-value in the permutation test, though assignments from big clones have predominantly more trustworthy (Suppl. Fig3E). Given the inherent difficulties in achieving confident assignments in single-GEM clones due to noisy pMHC and the inapplicability of smoothing, we recommend excluding high entropy pMHCs from TCR-pMHC assignments for single-GEM clones. This functionality is included in our workflow, as a parameter for assign_pmhc (Fig6B), to enhance the reliability of the analysis.

We recommend using both Rosner outlier detection test on smoothed data and permutation test on scaled data, thus addressing the question of whether a TCR is specific to one or more pMHCs within a given panel and if yes, is the observed pMHC signal in this clone is higher and more stable than simulated random, using a permutation test. The application of permutation test as a filtering step is optional, the decision whether to filter pMHC-TCR pairs is on the user, depending on the yield to rigor trade-off in the specific project. This functionality is included in filter_pmhc function (Fig6B,C bottom part), where a user can additionally filter responses by confidence score or average scaled UMI count within the clone. More details provided in the package documentation.

## Discussion

In this study, we have developed a computational pipeline, with statistical models to assign pMHC specificity to TCR clonotypes based on single-cell data of T-cells stained with barcoded pMHC multimers. ITRAP2 provides a robust strategy for TCR-pMHC pairing, even in datasets that are heavily confounded by different sources of noise. We applied a comprehensive denoising strategy, using the biological assumption of low expected pMHC UMI counts diversity, within one T-cell clonotype. We subsequently implemented an TCR-pMHC pairing approach, looking at the pMHC values distribution within a clonotype and across pMHC. Our method was successfully applied on both an in-house dataset including 8,141 GEMs generated from CD8⁺ T-cells stained with a library of 100 barcoded pMHC multimers across 11 donors and a 10x Genomics public dataset including 208,589 GEMs generated from CD8⁺ T-cell stained with 50 barcoded pMHC multimers, across 4 donors.

In the analyses, we observed that some pMHC multimers had particularly high background noise levels. These are mostly derived from certain A alleles, including HLA-A*03:01, HLA-A*11:01, and HLA-A*01:01 alleles. Given the high noise level, most TCRs were assigned to these HLAs when basing the analysis purely on the raw UMI count assignments. Applying the denoising approach developed here, largely resolves this issue. A deeper understanding of the intrinsic noisiness associated with specific HLA alleles in pMHC assays will be critical for optimizing the quality of data, and to obtain correct TCR-pMHC pairing, specifically related to the HLA types. The source of this noise remains unclear, potentially stemming from technical issues like cross-GEM contamination, low capture efficiency of pMHC associated barcodes, or biological factors such as unspecific and non-TCR mediated binding of pMHC multimers to T-cells. For example a KIR protein family, most notably KIR3DL2, had been shown to bind to HLA-A*11:01 and HLA-A*03:01. (Hansasuta et al. 2004)

Additionally, the size of the clone was found to largely impact the reliability of the pMHC assignment. However, the majority of clonotypes in most datasets would be small in size. Different immunological conditions are characterized by different levels of clonal expansion, for instance, autoimmune T-cell responses on average demonstrate less clonal expansion than acute viral infections or T-cell clones in cancer biopsies. Therefore, we would expect accurate TCR-pMHC assignments to be more challenging in autoimmune settings than in acute viral infections.

We additionally addressed the problem of the minimum panel size for single-cell TCR and pMHC assays. TCR-pMHC pairing statistically comes down to outlier detection, which is sensitive to the number of data points. After performing an in-silico simulation of smaller panel sizes, by removing random non-specific pMHC, we conclude that a panel of at least 25 pMHC is necessary for robust data analyses.

Given all the abovementioned, the advantage of our approach is the inclusion of multiple complementary metrics, such as the permutation test, the Rosner or extreme distribution test with their significance threshold (α), and the confidence score, which together provide flexible control over assignment stringency. Depending on the biological context - such as clone size, TCR-pMHC affinity, or cross-reactivity, users can adjust parameters and selection criteria to balance yield and precision. Importantly, we have designed our methodology integrated with the widely-used Seurat framework for single-cell analysis, ensuring that our method is accessible to a broader community. We expect that the approach can equally well be applied in CD4 T-cell detection strategies, using pMHC II multimers (Kocher et al. 2025).

As stated earlier, other tools including ITRAP1 and ICON, have previously been proposed for denoising and specificity assignment of single-cell pMHC TCR sequencing data both having certain specific shortcomings when it comes to signal normalization and identification of multiple targets for a given TCR. An important advantage of our approach over ITRAP1 is the capacity to adjust for pMHC-specific noise by using mean centering and SD scaling as part of the normalization process, as well as the use of a permutation test, that compares the measurements in the clone, with simulated random set of cells, that do not share the same TCR. In contrast, ITRAP1 only looks within the GEMs of a single clonotype and does not have an option to control the noise within pMHC. At the same time, ITRAP1 fails to assign multiple pMHC per clone, an important limitation, since TCRs often are not exclusive to a single pMHC (Wooldridge et al. 2012; Bentzen and Hadrup 2019). ICON relies on the presence of negative control multimers to simulate the background noise to normalize the pMHC UMI counts of interest. However, negative control can be challenging to identify across a broad range of HLA molecules, and when included, often represent a few data points only. ITRAP2 offers an advancement as it does not require negative controls, but rather uses the large pMHC data panel for background signal estimations.

The lack of a ground truth in pMHC-TCR pairing, makes it challenging to validate the ITRAP2 pipeline, based on existing data. Consequently, in our current assessment we rely on combinations of visual data assessments, the assumption of sequence similarities of TCRs recognizing the same pMHC, and comparisons with bulk T-cell screening data, where we only know the pMHC recognition per donor, not per TCR. The ultimate strategy for validation includes gene editing to express the identified TCRs in cell systems and experimentally validate their pMHC binding characteristics. This will provide ground truth of TCR specificity but remains a laborious and expensive technique.

A very critical step in the pMHC-TCR pairing approach is the TCR clonotype identification, which is not part of ITRAP2. Thus, it is the users responsibility to have robust clonotype identification established prior to specificity discovery. The clonotype labels will impact the smoothing and pMHC assignment drastically. The vast majority of T-cells have a unique pair of α and β chains (Brady et al. 2010), and for them clonotyping is trivial. However, different sources report from around 10% to 35% of human PBMC T-cells to have dual α chain mRNA expression (Zhu et al. 2023; Dupic et al. 2019), which can be inflated by ambient or doublet cell GEMs contamination in droplet based single-cell technology. Work from 1993 (Padovan et al. 1993) with co-staining of two Vα TCRs in mice, showed “up to one-third” cells containing dual alpha chains. Current TCR clonotyping strategies assign the alpha chain with highest transcript level as the ‘true alpha’, this may result in false assignments, and it has been demonstrated that the lower transcribed alpha, may just as well be responsible for the pMHC recognition, hence leading to indirect false pMHC-TCR pairing. Therefore accurate single-cell TCR clonotyping, that takes this into account is highly recommended, and improvements in the clonotype assignment will further improve the downstream pMHC assignment.

Moreover, since our the denosing strategy assumes identical TCR sequence within a clonotype, we would not recommend using ITRAP2 with smoothing at its current stage to analyse single-cell B-cell receptor recognition readouts, since more diversity within a clone is expected for B-cells, due to somatic hypermutation.

## Materials and methods

### DNA barcodes

The design of scRNAseq barcodes for 10x 5’ v2 has been described previously (Povlsen et al 2023). In brief, oligonucleotides with biotin attached were bought from LGC Biosearch Technologies (Denmark). These barcodes were dissolved to 100 µM in RNase-free water, while working stocks were further dissolved to 0.54 µM or 2.17 µM in PBS + 0.5% BSA + 1 mg/mL herring DNA + 2 mM EDTA. All DNA barcodes were stored at -20C.

### DNA-labelled pMHC multimers

DNA-labeled pMHC multimers were built as previously described (Bentzen AK et al 2016). In brief, 0.54 µM DNA barcodes compatible with capture in the 5’ v2 10x platform were attached to PE- or APC-conjugated dextran backbones (Povlsen et al. 2023). Then, 200 µM peptides and 100 µg/mL p*MHC (MHC with UV-cleavable peptide) were mixed in a 1:1 ratio, followed by exposure to 366 nm UV for 60 minutes at room temperature (UV exchange REF). The exchanged monomers were centrifuged at 3,300 g for 5 minutes at 4 C and added onto the DNA-labelled dextran backbone. This was followed by a 30-minute incubation on ice. Next, a barcode freezing buffer was added to reach a final concentration of: 1.5 µM D-Biotin, 0.1 mg/ml Herring-DNA, 0.5% BSA, 2 mM EDTA, and 5% glycerol. This was followed by incubation for 20 minutes, whereafter it was stored at -20C until use.

### Staining with barcoded pMHC multimers, sorting and single-cell capture of CD8⁺ T-cells - experimental part

Cryopreserved samples were thawed in 10 mL 37C RPMI+GlutaMax with 10% fetal calf serum (FCS) (R10). Samples were centrifuged at 1500 RPM for 5 minutes, followed by resuspension in 10 mL cold R10. Then, samples were centrifuged and resuspended in PBS + 0.5% BSA. Concurrently, 1.5 µL of each pMHC multimer was collected per patient and concentrated using Vivaspin 6 or 20 (100,000 MWCO, Sartorius, VS0602) to a volume of 60-70 µL. This was followed by two centrifugation steps of 10,000g at 2 minutes to sediment any remaining aggregates. The samples were centrifuged and the upconcentrated multimer pool was added to each sample and incubated for 60 minutes at 4C. Next, samples were washed twice in PBS + 0.5% BSA, followed by adding 2.5 µL Human TruStain FcX Fc blocking reagent and incubated for 10 minutes at 4C. TotalSeq-C hashtag antibodies (BioLegend, 422302) were added, which was followed by a 15-minute incubation period. Next a 5× antibody mix composed of CD8-BV480 (BD566121, cloneRPA-T8) (final dilution 1/50), dump channel antibodies: CD4-FITC (BD 345768) (final dilution 1/80), CD14-FITC (BD 345784) (final dilution 1/32), CD19-FITC (BD 345776) (final dilution 1/16), CD40-FITC (Serotech MCA1590F) (final dilution 1/40), CD16-FITC (BD 335035) (final dilution 1/64) and a dead cell marker (LIVE/DEAD Fixable Near-IR; Invitrogen L10119) (final dilution 1/1000) was added to the samples and incubated for 30 minutes at 4 C. Samples were washed three times in PBS + 0.5% BSA before acquiring on a BD Aria Fusion cell sorter. 2,500 pMHC multimer+ and CD8⁺+ cells from each donor were sorted collectively into a BSA-saturated 1.5 mL eppendorf tube with 100 µL PBS+0.5% BSA. A total of 28,000 cells were sorted, and assuming a transfer of 60% of the sorted cells, we loaded 17,000 cells onto a Chromium Controller. We utilized the 10x Genomics 5’ v2 chemistry to capture gene expression, V(D)J, pMHC barcodes, and hashtag-associated barcodes. The following downstream process was carried out according to the manufacturer’s protocol (Chromium Next GEM Single Cell 5’ Reagents Kit v2, CG000330). The product was sequenced on a NovaSeq 6000 (NovoGene, UK).

#### Bioinformatics

Processing of 10x single-cell data

Hashing and peptide-MHC barcode, gene expression, and TCR enriched reads were provided in fastq format and were processed using 10x Genomics Cellranger multi v7.0.0(“V(D)J Cell Calling Algorithm -Software -Single Cell Immune Profiling -Official 10x Genomics Support,” n.d.)(10x Genomics, n.d.-b)

### Postprocessing of 10x hashing

The donors were demultiplexed using the Seurat functionality. HTO UMI count data was normalized using NormalizeData function with normalization.method = “CLR” parameter. Donors were labeled using the HTODemux function, with default parameters(10x Genomics, n.d.-b, “Integrated Analysis of Multimodal Single-Cell Data” 2021).

### Postprocessing 10x TCR data and clonotyping

TCR-seq outputs from cellranger multi filtered_contigs.fasta and filtered_contig_annotation.csv were further processed with change-o ver 1.2.0 scripts from the immcantation pipeline(Gupta et al. 2015), with the following workflow: AssignGenes.py, MakeDb.py, ParseDb.py, DefineClones.py. Only clonal assignments with UMI count higher than 2 supporting each TCR chain were included. Clonotypes with only beta chain and missing α chain were also included in the analysis.

### Quality control

We filtered GEMs that have higher than 10% mitochondrial transcripts, and lower than 300 genes expressed. Also, gems with a number of GEX UMI counts higher than 4000 were excluded. Specifically to compare with ITRAP1, the 10x Genomics dataset was additionally processed with a 20% mitochondrial transcripts cut off to match previous data input.

### Data denoising

We use the Seurat framework to apply the pMHC denoising pipeline. We recommend storing raw counts in object@assays$pMHC@counts.

UMI counts scaling is performed in the following way. We mean-center and standard deviation scale each pMHC across all donors

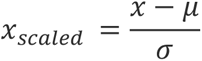

Next, we remove every outlier, defined as |𝑥_𝑠𝑐𝑎𝑙𝑒𝑑_ |>3, and calculate the mean and standard deviation again until a pMHC vector is left with no outliers, then these mean and standard deviation are used to calculate a final pMHC UMI z-score vector. In case the mean or sd becomes zero after the kth iteration, then the k-1th value is taken. We could not use median-based parameters, single sparse single-cell readouts end up in most medians and the median deviation being zero. This functionality is implemented in the Seurat-applicable method ScaleDataNoOutliers. The Z-score matrix will be stored in object@assays$pMHC@data

### Data Preparation

For each selected clone, based on a predefined size threshold (default=3), peptide-MHC (pMHC) count data is recommended to be extracted for analysis, after scaling, from object@assays$pMHC@data layer.

### Noise assessment

Each pMHC’s noisiness level is assessed via entropy metrics. Entropy, a concept from information theory, quantifies the level of uncertainty or randomness in the pMHC count values across different clonotypes. In this study, we utilize entropy to measure the “noisiness” of the pMHC data, allowing for an assessment of informative pMHC. As a very limited set of pMHCs are expected to be recognized within a given T-cell clone, a higher entropy value indicates greater variability in pMHC count values, suggesting less reliability in the signal, while lower values indicate less noisy data.

The entropy H is calculated as follows:

1. We calculate quantiles for each pMHC (.2, .4, .6, .8, 1 by default) across all gems.
2. Quantiles of the pMHC count levels are determined, and if there are fewer than two unique quantiles, the entropy is set to 0.
3. The pMHC count values for a given GEM are binned according to these quantiles
4. Finally, the entropy is calculated using the formula:

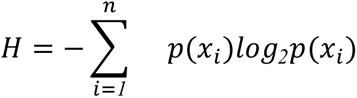

- 𝐻 — entropy
- 𝑛 — total number of unique quantiles
- 𝑝(𝑥_𝑖_) — probability of observing i-th pMHC UMI count quantile

H is calculated within each clonotype, and the average is taken for the whole pMHC.

### Application of LOESS Smoothing

The LOESS smoothing was implemented using the loess function in R, following these steps:

1. Clone and pMHC Selection:

Clones exceeding a specific size threshold (default 3) were identified for LOESS smoothing. For each of these clones, a pMHC z-scores matrix is extracted, which is stored in object@assays$pmhc@data slot.

1. Smoothing Process:

For each pMHC vector within a clone, a LOESS model was fitted to the sequential order of counts. Model parameters - span, degree, and family are flexible, can be provided as a parameter in smooth_pmhc:

- span: the fraction of data points used in the smoothing process. The default is 1 for maximum smoothing.

- degree: the degree of polynomial used. (default is 1)

- family: the type of response distribution to adapt the smoothing approach. We recommend choosing “symmetrical”, due to its less sensitive effect on outliers.

The loess() function was applied within assign_pmhc() to model and predict() to smooth the counts.

It’s only possible to smooth scaled pMHC counts within clones of at least size 3, the rest of the clonotypes are left unsmoothed and copied from the scaled matrix, and the user can choose to still proceed with TCR assignment on scaled counts for them.

This functionality is Seurat-friendly and implemented as a method called smooth_pmhc. The smoothed matrix will be stored in object@assays$pmhc@scale.data.

### pMHC-TCR assignment

The next step is an iteration over clone, where smoothed pMHC UMI z-scores are TCR clone-based pseudobulked as average values. We detect outliers within each clone and assign detected as outliers by either Rosner or Extreme distribution tests pMHC as specificities.

#### Rosner test

The collapsed pseudo-bulk smoothed and scaled pMHC UMI counts are tested to have outliers by the Rosner outlier test. It’s a multiple outlier detector, which can help us identify cross-reactive clones. It has an option to detect k (up to 10) outliers, which is suitable since we don’t expect more than 10 reactivities per clone. By default, the test calculates the statistics based on mean and standard deviation, re-estimating them each k^th^ time, without the most extreme data point. We introduced an extra option, that we believe is more suitable for our purposes, by removing x (3 by default) most extreme values and estimating mean and standard deviation once.

We use rosnerTest() R function from the EnvStats(Millard 2013) function with an extra option to choose the stable mean and standard deviation estimation, without the most x extreme values (default 3) to calculate a test statistic. By default, the Rosner test does not output a p-value but calculates a test statistic R that is normalized by predefined α, depending on the user’s preferred rigor. Since R is a z-score-like statistic, centered around zero, we used normal distribution with mean=0 and sd=1 using rnorm() R function to calculate the p-value for a given pMHC.

#### Extreme distribution test

We observed that pseudo-bulked, smoothed and scaled pMHC values within a clonotype follow extreme distribution, the type of statistical distribution with a heavy tail and one-sided peak. (Fig2D) We fit an extreme distribution model to each clonotype, estimating key parameters such as scale (a measure of variability), location (a metric representing the peak or mode in case of Gumbel distribution), and shape (a parameter that defines the tail behavior)(Gilleland and Katz 2016). We used a specific form of generalized extreme distribution, called Gumbel distribution(Gumbel 2012), which is particularly suited for our data since it models the maximum values well and assumes a right-sided tail. This distribution has a fixed shape parameter, simplifying our modeling as it only varies by location and scale. The location parameter typically approaches zero, which fits our data where extreme pMHC values are not exceedingly high. For each pseudobulk pMHC value, we calculate the probability that it fits within the extreme distribution with the estimated parameters. We classify pMHC values as outliers if the calculated probability (p-value) falls below a predefined α, assigning these outliers to the respective clone. The Gumbel distribution also tends to have smaller tails compared to other extreme value distributions. This results in generally higher scale values, leading to more stringent criteria for identifying outliers in TCR-pMHC assignments.

We use fevd() R function from extRemes to fit an extreme distribution for pseudobulked pMHC values, with the type parameter ‘Gumbel’. Given the extracted location, shape, and scale parameters, we use pevd() R function to calculate the probability of having a specific value for each pMHC within a clone. The ones that are lower than the set α level will be assigned a specificity

This functionality is implemented as a Seurat method called assign_pmhc, with the parameter “assignment” that determines the TCR-pMHC pairing option. The default method for the assignment is chosen to be Rosner.

### Permutation test

Each pMHC-TCR pair that went through the pMHC-TCR assignment procedure is subjected to a permutation test. N (default 10000) pseudoclones (groups of cells that don’t share the same TCR, to simulate random signal) are generated with the same size as the analyzed clonotype, and we calculate the proportion of pseudoclones, whose average z-score and percent of gems, with z>0.1(GEMs having a rescaled pMHC count above 0.1, is a proxy for a non-zero raw UMI count). analyzed pMHC z-scores are equal or greater than the observed one. Two resulting p-values are united into a single probability with the Fisher method. This functionality is implemented within the assign_pmhc method.

### Confidence scoring

Each pMHC-TCR pair that underwent the pMHC-TCR assignment procedure is subjected to confidence scoring. This functionality is implemented within the assign_pmhc method.𝐶𝑜𝑛𝑓𝑖𝑑𝑒𝑛𝑐𝑒 = 𝛼𝐶 + 𝛽𝑙𝑜𝑔*_10_* (𝑆 + *1*) + 𝛿𝑑 − 𝛾𝐸

- 𝛼, 𝛽, 𝛾 — weights
- 𝐶 — clonotype concordance
- 𝑆 — clone size
- 𝐷 — difference between mean scaled pseudobulked pMHC UMI counts of specific mean pseudobulked background pMHC
- 𝐸 — pMHC entropy

### Validation with bulk pMHC barcoding

#### Adjusted estimation of pMHC specific cells in bulk screen

The adjusted estimated number of pMHC-specific cells captured by the bulk assay ( 𝑁_𝑖_^𝑏𝑢𝑙𝑘,𝑎𝑑𝑗^) was calculated by scaling the bulk-estimated frequency of each pMHC response by the total estimated frequencies of all pMHC responses, multiplied by number of pMHC-specific GEMs in single-cell assay. Specifically, for pMHC

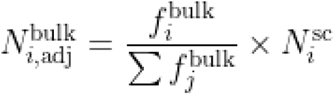

- 𝑁^𝑠𝑐^ _𝑖_ - number of pMHC specific T-cells across every clone, measured by single-cell
- 𝑓_𝑖_ - estimated T-cell frequency for a given pMHC
- 𝑓_𝑗_ - vector of T-cell frequencies for a given sample

For the Pearson correlation coefficient, calculated between adjusted estimated pMHC specific T-cell number and single-cell calculated pMHC specific GEMs number (Fig4A), only bulk responses with estimated frequency above 1% were taken into account.

#### Precision and Sensitivity calculation

To calculate the accuracy of the assignment we used a bulk pMHC screen of the same healthy donors with the same pMHC panel, where we know per donor pMHC response and approximate fraction of antigen-specific T-cells in the sample. Precision was calculated as a ratio of true positive responses (TCR-pMHC pair assigned within a given donor with the same pMHC response detected in bulk) to a number of all TCR-pMHC assignments. Sensitivity was defined as the ratio of a number of true positives to a sum of true positives and true negatives (defined as a pMHC response detected in bulk and not detected in single-cell).

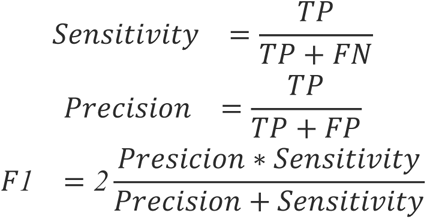

- TP (True Positive) — number pMHC responses present in the donor in single-cell and in bulk
- FN (False Negative) — number of identified pMHC responses in the bulk but not detected in single-cell
- FP (False Positive) — number pMHC responses identified in a donor in single-cell but not in bulk

### Intra vs Inter similarity calculation

To calculate intra and inter-similarities, we scored all TCRs assigned to a given pMHC against the same set of TCR (excluding self) and recorded for each TCR, the score of the most similar hit (intra-similarity). Likewise, the TCRs were scored against a random subset of size equal to the inter-similarity analysis of TCRs with assigned specificities to any different pMHC and the most similar score was recorded (inter-similarity). In both cases, the similarity between two TCRs was calculated from the summed score of the pairwise α- and β-chain similarities calculated using the TCRbase scoring model. We conducted this intra versus inter similarity evaluation for the 10x data set, annotated by the three methods. Here, the notations for ICON and ITRAP1 were obtained from(Montemurro et al. 2023), and the annotation of ITRAP2 was made using the Rosner test with a permutation p-value threshold of 0.01

### Optimal panel size

In order to determine the minimal theoretical number of barcoded pMHC multimers in single-cell experiments for good-quality TCR-pMHC pairing we performed the following analysis. Clones with high confidence scores (> .5 for 1OS dataset and >2 for 10x dataset, different cut-offs due to big differences in dataset sizes) were selected, which means every clone has at least 1 specificity with a good level of certainty. We would perform the Rosner test to assign the specificity within a clone with k (expected number of outliers) equal to the previously paired number of pMHC multimers, repeating the original procedure x times, each iteration removing 1 random multimer that wasn’t paired before, with a full panel. We stop when the Rosner test doesn’t identify the expected multimer as an outlier, save a number of iterations, this number of iterations is interpreted as a panel size where we lose sensitivity. We also differentiate detection error from Rosner test errors (Fig5F), since EnvStats implementation of the test raises error when the provided vector has almost no variance, or k is greater than n-2, where n is a number of non-missing measurements.

### Code and data availability

Our in-house generated dataset is available on ArrayExpress with the accession number E-MTAB-13758.

10x data for 4 separate donors is available here(10x Genomics, n.d.-a). https://www.10xgenomics.com/datasets/cd-8-plus-t-cells-of-healthy-donor-1-1-standard-3-0-2 https://www.10xgenomics.com/datasets/cd-8-plus-t-cells-of-healthy-donor-2-1-standard-3-0-2 https://www.10xgenomics.com/datasets/cd-8-plus-t-cells-of-healthy-donor-3-1-standard-3-0-2 https://www.10xgenomics.com/datasets/cd-8-plus-t-cells-of-healthy-donor-4-1-standard-3-0-2

The R package with documentation is available through the github page https://github.com/SRHgroup/ITRAP2

## Funding

This study was funded by the Danish Cancer Society, grant no. R302-A17640 and the European Union’s Horizon 2020 research and innovation program under ImmunoSABR grant agreement no. 733008 to SRH and MK; The European Research Council, ERC CoG project MIMIC (101045517) to SRH, the Novo Nordisk Foundation, Data Science Collaborative Research Programme - 2024, grant no. 0089619 (DIRM) to MN and SRH, the Lundbeck Foundation grants (R324-2019-1671 and R322-2019-2445) to AKB. MK received funding from the European Union’s Horizon 2020 research and innovation program under the Marie Sklodowska-Curie grant agreement no. 713683 (COFUNDfellowsDTU)

## Supporting information

Supplementary_table_1_10x_TCR-pMHC_ITRAP2

Supplementary_table_2_in-house_TCR-pMHC_ITRAP2

Supplementary_table_2_in-house_TCR-pMHC_ITRAP2

Supplementary_table_3_10x_TCR-pMHC_ITRAP1

Supplementary_table_4_10x_dataset_donor_HLA_typing

Supplementary_table_5_in-house_dataset_donor_HLA_typing

Supplementary_table_6_bulk_screening

## Conflict of interest

GN, MV, KJ, JN, NK, LG, LV, MK, MN declared no competing interests. AB, SH AKB and SRH are co-inventors on a patent covering the use of DNA barcode-labeled MHC multimers (WO2015185067 and WO2015188839), which is licensed to Immudex

---

**Supplementary Figure 1.**
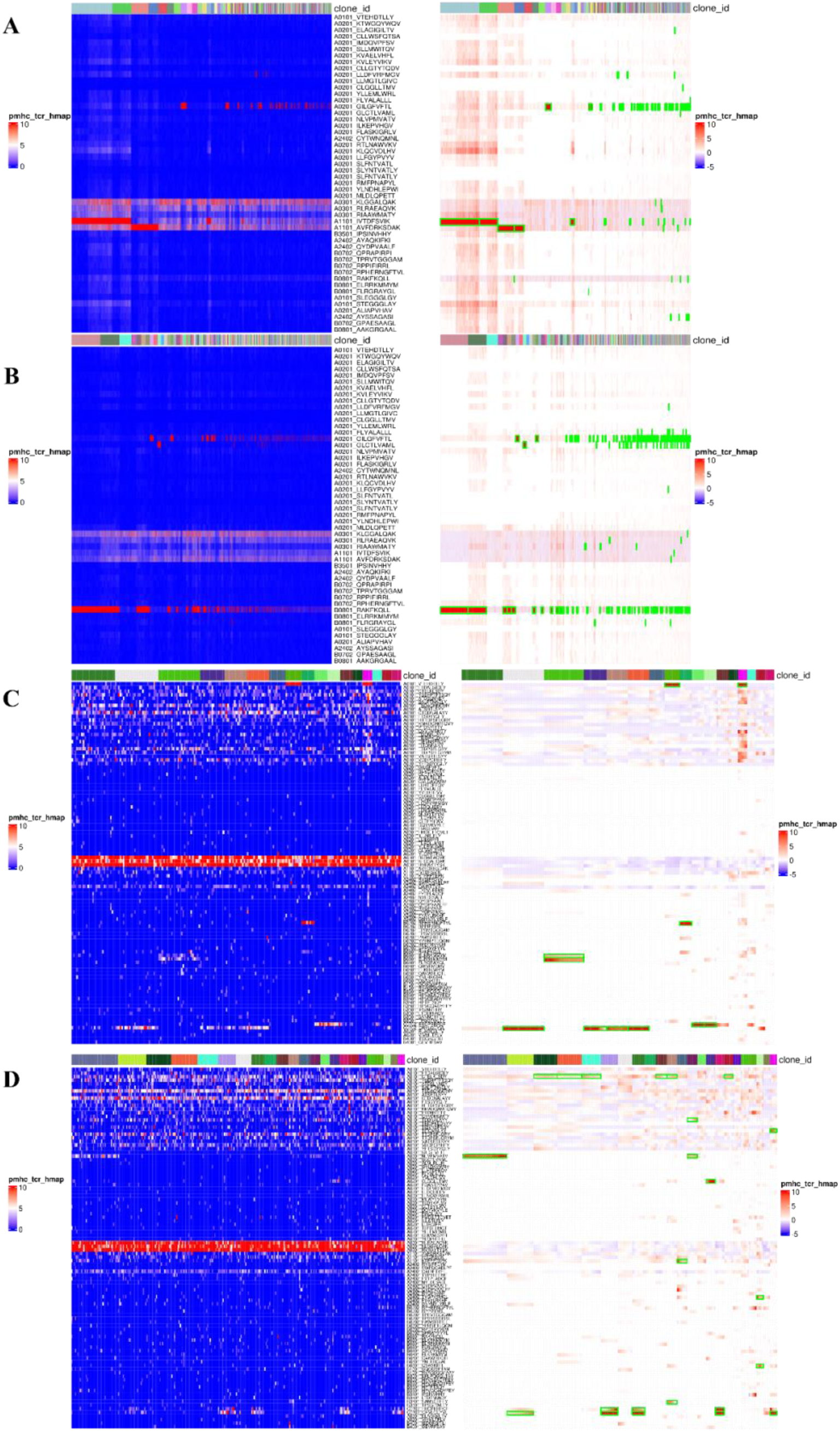
**A)** Heatmap with raw UMI counts grouped by clonotype on the left, and with smoothed counts with highlighted specificities on the right, for donor #1 in the 10x dataset. **B)** Same as Suppl Fig1A but for donor #2. **C)** Same as Suppl Fig1A but donor BC360 in the in-house dataset. **D)** Same as Supl Fig1A but donor BC360 in the in-house dataset.

**Supplementary Figure 2.**
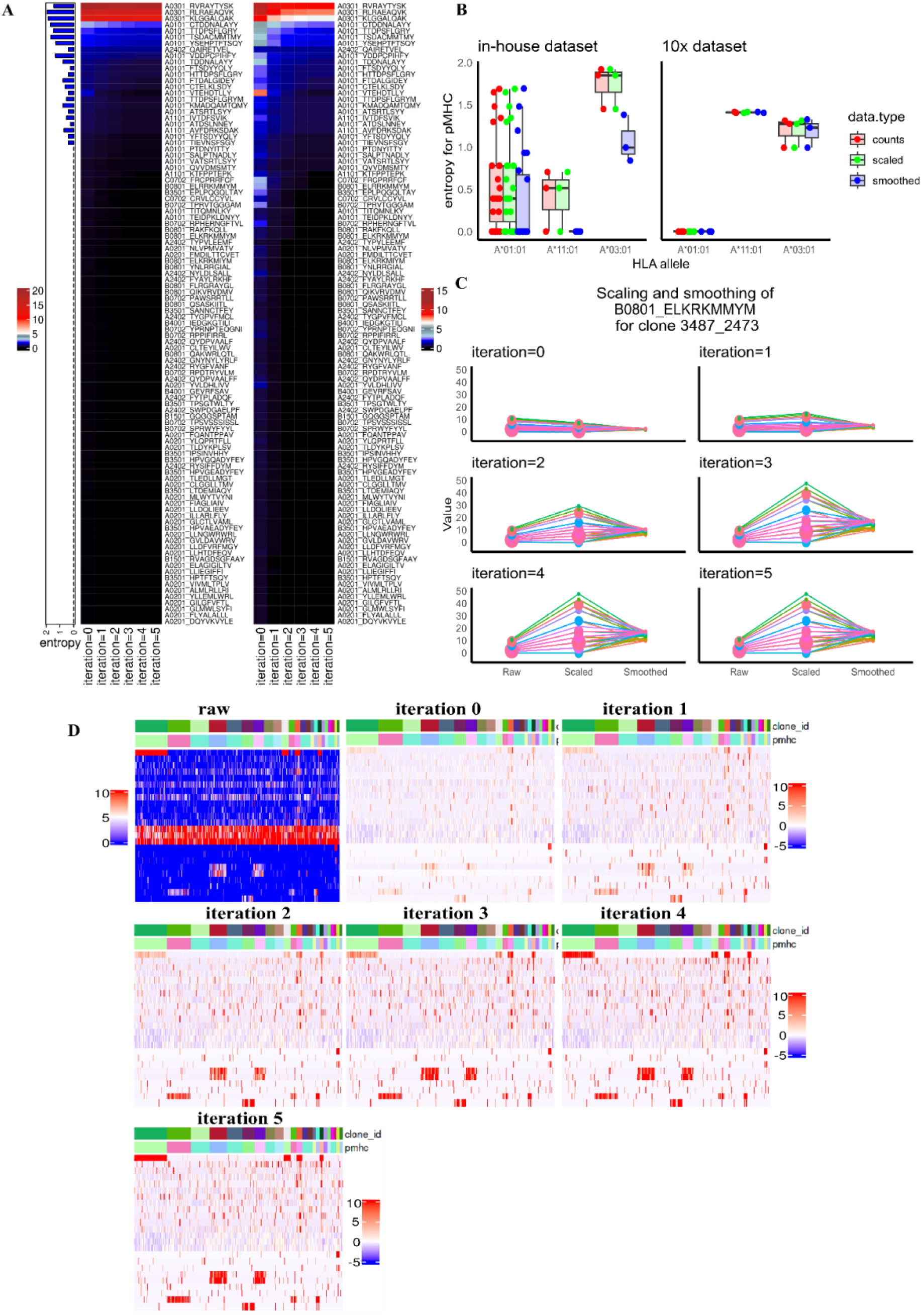
**A)** Heatmap means on the left and standard deviation on the right for each pMHC in the in-house dataset, with annotated noisiness levels (entropies) on the left barplot. Each column on the heatmap represents re-estimated means and standard deviations after one iteration of outlier removal. **B)** Boxplot showing noisiness level (entropy) changes for intrinsically noisy pMHCs for each normalization step. **C)** Changes in z-scores and resulting smoothed pMHC values from raw counts for a single clone when means and standard deviations are estimated at every step of outlier removal. **D)** Raw and then scaled pMHC UMI count heatmaps, with z-scores calculated using means and standard deviations from each outlier removal iteration.

**Supplementary Figure 3.**
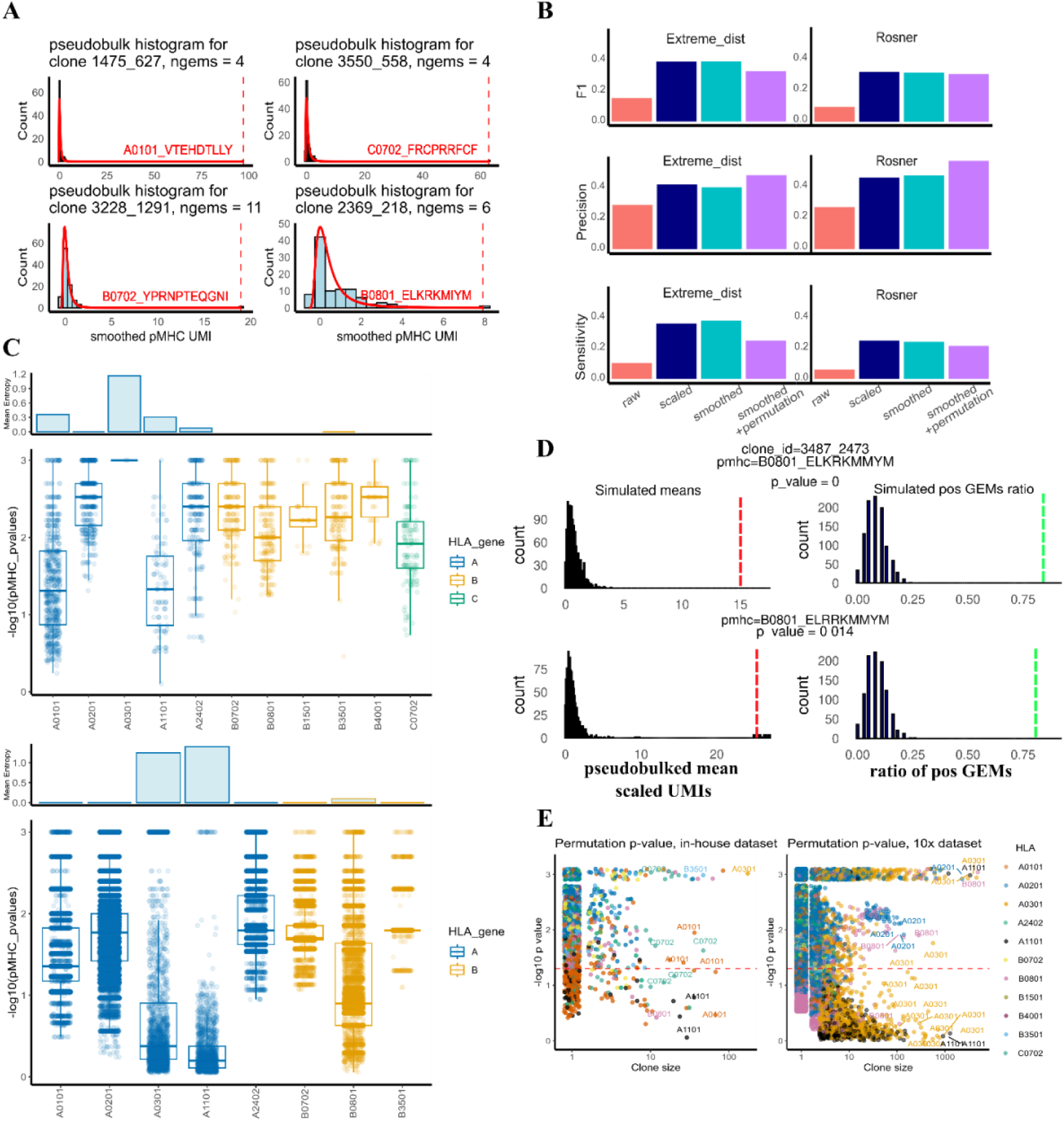
**A)** The observed pMHC UMI count distribution shown as a histogram, overlaid with the density plot calculated from a simulated extreme distribution using parameters estimated from the observed distribution. **B)** P-values for the permutation test displayed next to the smoothed pMHC UMI count distributions, demonstrating significance where expected. **C)** Boxplots showing permutation p-values per pMHC grouped by HLA alleles (in-house dataset on the left, 10x dataset on the right). **D)** Relationship between permutation test p-value and clone size.

**Supplementary Figure 4.**
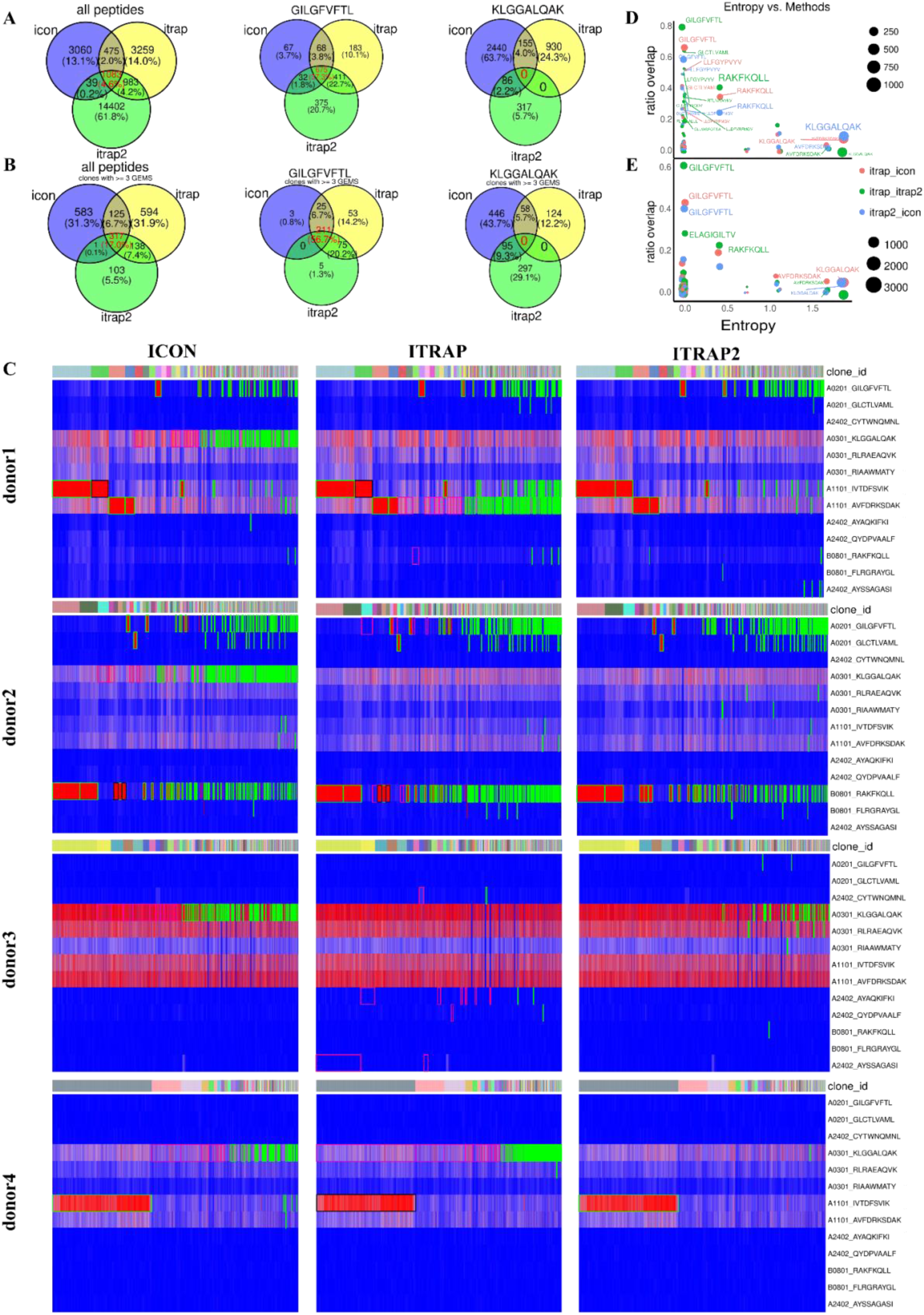
**A)** Overall pMHC–TCR pair assignment overlap between the three methods, shown for a low-noise peptide (GIL) and a high-noise peptide (KGL). **B)** Same as Supl Fig4A but restricted to clones with ≥3 GEMS. **C)** Relationship between noise level and agreement in pairing between different methods. **D)** Heatmaps with green frames highlighting assignments from each method; black frames marking visually apparent false negatives from ICON or ITRAP1; and pink frames marking visually apparent false positives.

**Supplementary Figure 5.**
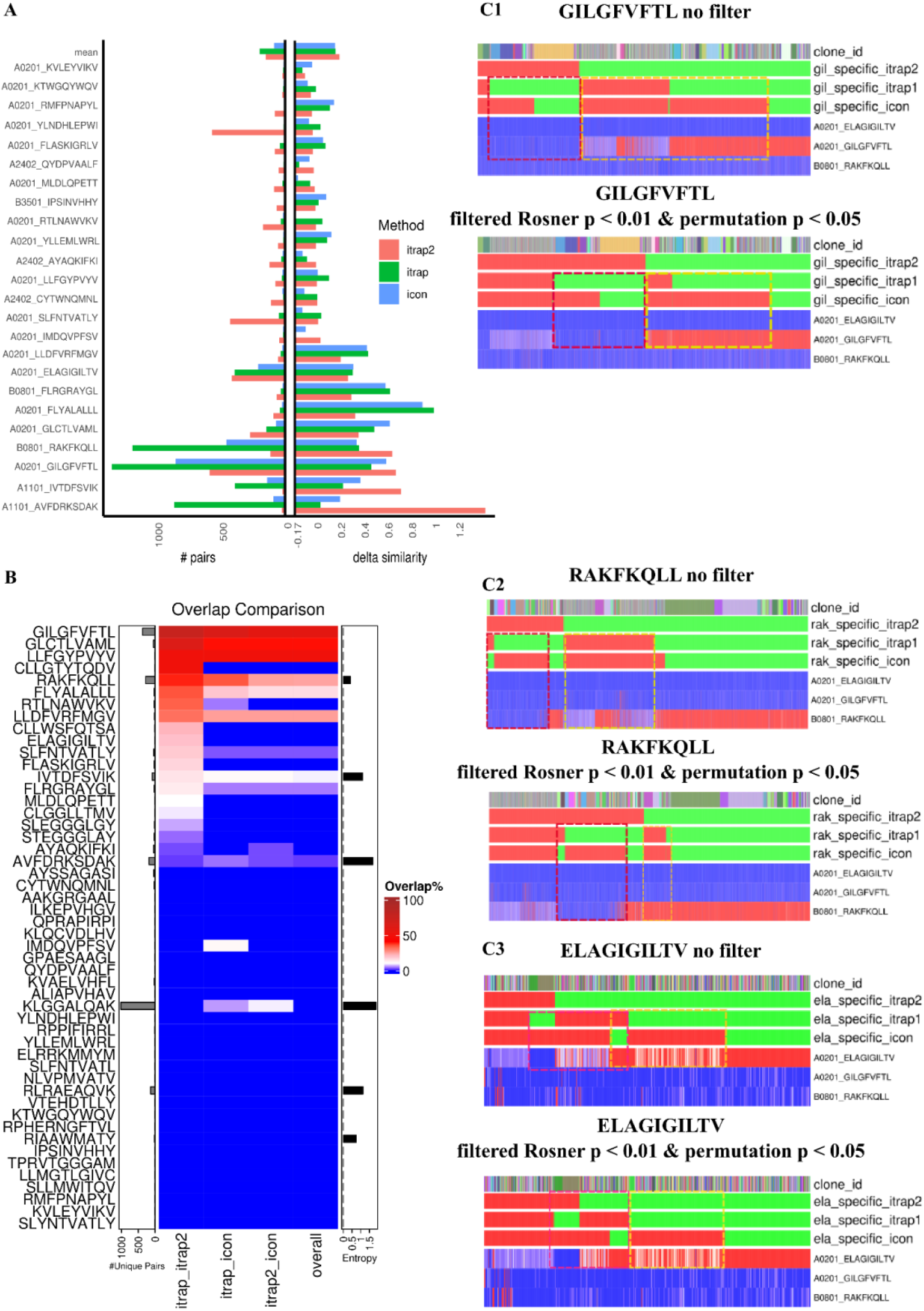
**A)** The left barplot shows the total number of detected TCR–pMHC pairs per pMHC; the right barplot shows delta similarity, i.e., the difference in TCR sequence similarity within each pMHC compared to non-specific TCRs. **B)** Heatmap showing overlap between methods per column for different pMHCs (rows), with entropy annotated on the left barplot. **C)** Heatmaps for the three peptides producing the most TCR–pMHC pairs with the least noise—GILGFVFTL (C1), RAKFKQLL (C2), ELAGIGILTV (C3). Each heatmap includes method-specific annotations indicating whether a clonotype is classified as positive or negative. For ITRAP2, the Rosner test annotations are shown in the upper heatmap and the permutation test in the lower one. Yellow dotted marks indicate suspected false negatives from ICON or ITRAP1 where ITRAP2 classified correctly; purple marks indicate suspected false positives from ICON or ITRAP1 where ITRAP2 classified correctly.

## Notes

https://www.ebi.ac.uk/biostudies/arrayexpress/studies/E-MTAB-13758

### Summary of Updates

Removed duplicated figures, previously added duplicated figures by mistake

